# An oxidative mutational signature in normal tissues of a monoallelic *MUTYH* carrier

**DOI:** 10.64898/2026.11.12.766009

**Authors:** Christopher M. Grochowski, Yang Zhang, Eric Venner, Marie-Claude Gingras, Sravya Bhamidipati, Ziad Khan, Jesse Farek, Luis F. Paulin, Yilei Fu, Chenghang Zong, Fritz J. Sedlazeck, Harsha Doddapaneni, The SMaHT Consortium, Richard A. Gibbs

**Author notes:** Correspondence: Richard A. Gibbs, Human Genome Sequencing Center, Baylor College of Medicine, Houston, TX, USA.

## Abstract

Biallelic loss of *MUTYH* causes *MUTYH*-associated polyposis (OMIM: 608456), characterized by the oxidative base-excision-repair mutational signature SBS18, but whether the roughly 1 in 100 individuals who carry a single pathogenic *MUTYH* allele acquire a mutational phenotype in normal tissue is unknown. Analysis of multi-tissue sequencing of 25 donors from the Somatic Mosaicism across Human Tissues (SMaHT) Network identified three heterozygous carriers of *MUTYH* p.Gly396Asp. One carrier showed SBS18 across multiple normal tissues, ranking first of 25 donors with three independent methods and remaining significant after adjustment for tissue, without a somatic, epigenetic or transcriptional second hit at the *MUTYH* locus. A single hypomorphic *MUTYH* allele can therefore leave a measurable oxidative mutational mark in healthy tissues, supporting tissue-level mutational readouts as an approach to cancer risk stratification within this common carrier state.

## Main text

Inherited defects in DNA damage repair (DDR) genes are among the most actionable forms of cancer predisposition, yet almost everything known about how pathogenic alleles drive tumorigenesis comes from studies of advanced disease^1^. This bias complicates mechanism-based early prediction of disease risk, particularly in carriers who have only one copy of a pathogenic allele for recessive conditions that have only a modest increase in cancer rates, established from population studies.

Reactive oxygen species modify the DNA base guanine into 8-oxoguanine (8-oxoG) which can pair with adenine, instead of the expected cytosine. The mis-incorporated adenine is normally excised by a *MUTYH* encoded glycosylase, however biallelic pathogenic variants in *MUTYH* disrupt this process, resulting in frequent G:C>T:A transversions. Further, biallelic pathogenic variants in *MUTYH* frequently result in *MUTYH*-associated polyposis (MAP; OMIM: 608456)^2,3^, leading to cancer in approximately 70% of individuals without surveillance^4^. Tumors and normal cells from these individuals with biallelic pathogenic *MUTYH* variants and G:C>T:A transversions exhibit the characteristic single-base substitution (SBS) mutational signatures, SBS18 and SBS36 signatures^5–8^.

Heterozygous carriers of pathogenic *MUTYH* variants, about 1.3% of individuals globally^9^, have a modest overall increase in colorectal cancer risk (odds ratio 1.1–1.2)^10–12^. When SBS18 has been found in tumors from carriers it has usually been attributed to somatic loss of the wild-type allele^13,14^, although a recent tumor-only analysis proposed that haploinsufficiency alone can raise SBS18^15^. Whether a single pathogenic *MUTYH* allele can produce a characteristic mutational signature in non-neoplastic tissue has not been tested.

We analyzed DNA sequence data from multiple tissues from 25 donors (the P25 cohort) studied by the Somatic Mosaicism across Human Tissues (SMaHT) network^16^. Germline variants in DDR genes identified from sequencing of DNA extracted from blood tissue included a heterozygous pathogenic allele, *MUTYH* c.1187G>A (p.Gly396Asp) in each of three donors (SMHT020, SMHT031, SMHT043). This unexpectedly high frequency (3/25) relative to reported population data^17^ prompted additional validation that confirmed their presence (Fig. 1a,b).

**Fig. 1.**
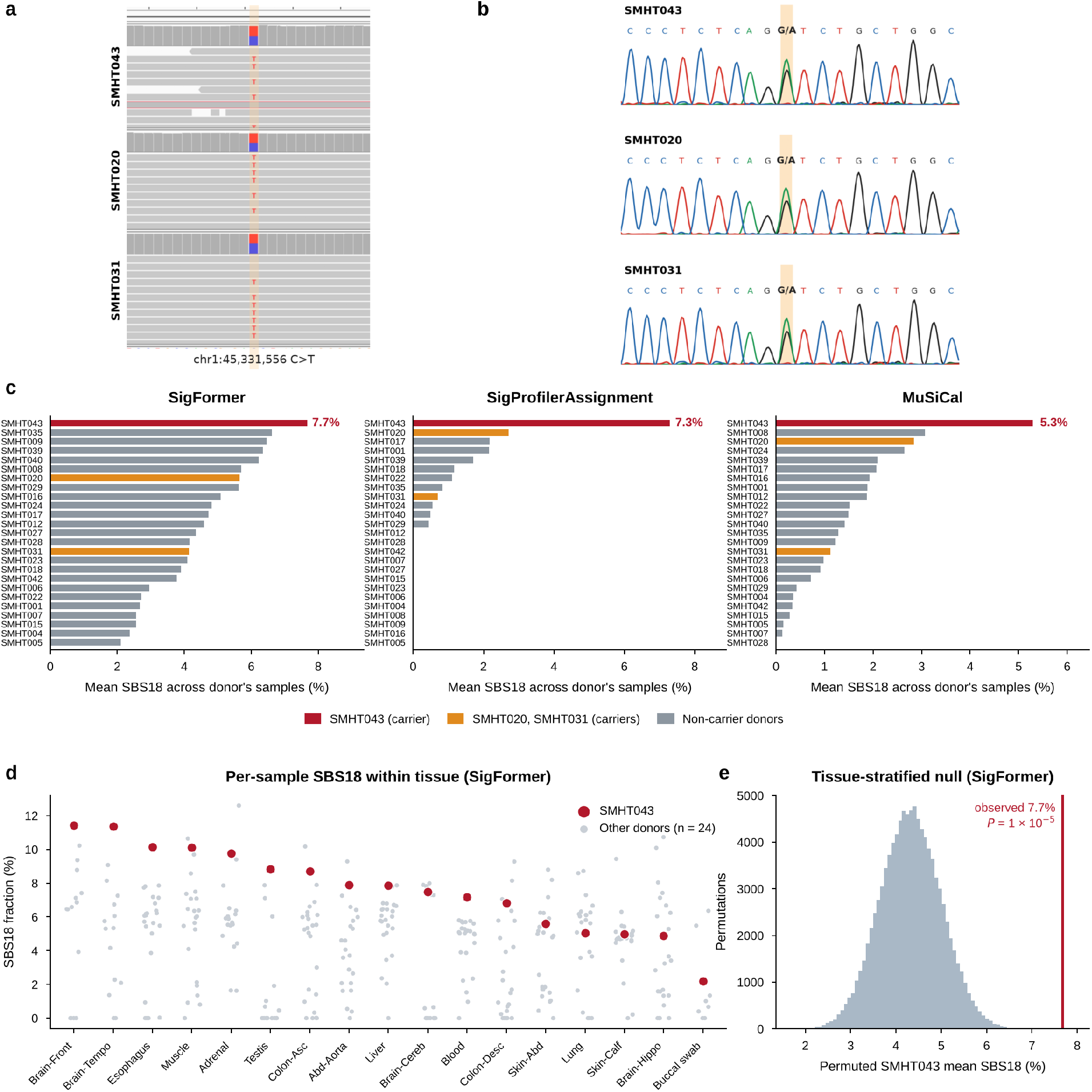
A heterozygous *MUTYH* p.Gly396Asp allele in three SMaHT donors and an oxidative mutational signature in one. **a**, Whole-blood short-read whole-genome sequencing reads at chr1:45,331,556 (GRCh38) for SMHT043, SMHT020 and SMHT031 (top to bottom); the alternate T corresponds to c.1187G>A on the *MUTYH* coding strand. **b**, Sanger sequencing of the coding strand in the same donors showing the heterozygous G/A peak (highlighted). **c**, Donor-level mean SBS18 fraction across all tissues of each of the 25 P25 donors with three methods (SigFormer, SigProfilerAssignment, MuSiCal; SmahtSNV 2.0.0 PASS calls, 381 samples); SMHT043 (red) ranks first with each; the other two carriers (SMHT020, SMHT031) are shown in orange. **d**, Per-sample SBS18 fraction (SigFormer) within each tissue, ordered by the SMHT043 value (red, SMHT043; grey, other donors). **e**, Null distribution of the SMHT043 donor-level mean SBS18 fraction from 100,000 tissue-stratified donor-label permutations of the SigFormer attributions; red line, observed value 0.077; upper-tail *P* = 1 x 10^−5^.

We next used three independent methods to identify patterns of somatic single-nucleotide variants (sSNVs) in all 381 tissue data sets from the 25 SMaHT donors (SmahtSNV 2.0.0 consensus calls, PASS filter; 536,748 sSNVs; Methods). Comparison with data from the Catalogue of Somatic Mutations in Cancer (COSMIC) database (v3.4) showed that SMHT043 had the highest donor-level mean SBS18 of all 25 donors with every method (SigFormer 7.7%, SigProfilerAssignment^18^ 7.3%, MuSiCal^19^ 5.3%) (Fig. 1c). Six SMHT043 tissues exceeded 5% SBS18 by both SigProfilerAssignment and MuSiCal: esophagus (24.8%/13.2%), skeletal muscle (24.4%/10.1%), temporal lobe (22.4%/15.3%), adrenal (21.3%/13.4%), frontal lobe (19.2%/10.4%) and blood (12.0%/5.2%); the first five were also the tissues with the highest SigFormer SBS18 fractions in SMHT043 (Fig. 1d). Neither of the other carriers (SMHT020 and SMHT031) showed cohort-level SBS18 elevation, and SBS36 was not consistently assigned across methods (Methods). The SMHT043 signal was not explained by tissue composition or mutation count: in a tissue-stratified donor-label permutation test (100,000 permutations; SigFormer attributions) the donor mean lay in the extreme upper tail of the tissue-matched null (observed 0.077; *P* = 1 x 10^−5^; Fig. 1e), and a tissue-adjusted beta-binomial model of SBS18-attributed over total substitutions gave an odds ratio of 1.96 (95% confidence interval (CI) 1.48-2.54; likelihood-ratio *P* = 1.9 x 10^−5^). The enrichment was reproduced with SigProfilerAssignment and MuSiCal (permutation *P* ≤ 9 x 10^−5^; odds ratios 19.1 and 5.0; Extended Data Fig. 1).

No somatic single-nucleotide or structural variant was found at the *MUTYH* locus in any tissue of the three carriers. Long-read CpG methylation profiles across *MUTYH* placed each carrier within the distribution of the other P25 samples of the same tissue class, with an unmethylated promoter/CpG island in every carrier tissue; the tissue-specific methylation of the locus, including lower gene-body 5-methylcytosine (5mC) in skeletal muscle and in brain, was shared with non-carriers (Fig. 2a and Extended Data Fig. 2). In bulk RNA-sequencing, the variant allele was detected in transcripts from every carrier tissue with RNA data, with no reduction in read coverage over downstream exons, and whole-gene *MUTYH* transcript abundance in all three carriers was within the cohort distribution, with no difference between carrier and non-carrier samples (Fig. 2b,c). The oxidative phenotype observed in SMHT043 therefore arises from a normally expressed, unsilenced locus at which both alleles are transcribed, which is the expected consequence of a missense allele that retains partial glycosylase activity (about half of wild-type 8-oxoG:A repair activity in a saturation variant-effect map^20^) while still leaving transcription of *MUTYH* intact.

**Fig. 2.**
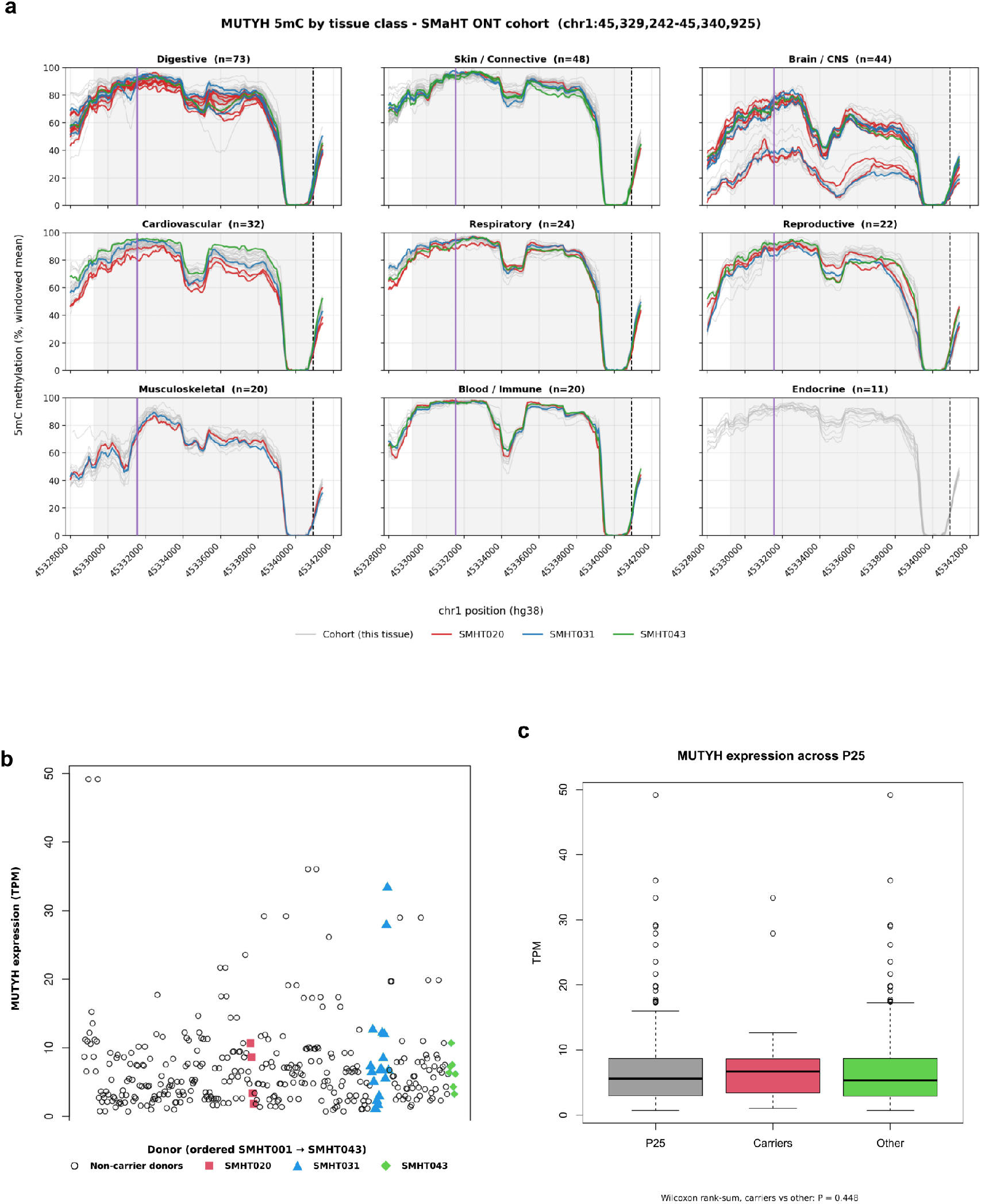
No epigenetic or transcriptional second hit at *MUTYH* in carriers. **a**, Oxford Nanopore 5-methylcytosine (5mC) across *MUTYH* (shaded; chr1:45,329,242–45,340,925, GRCh38) shown as the windowed mean percentage of methylated reads per sample, by tissue. Color lines, the three carriers (SMHT020, red; SMHT031, blue; SMHT043, green); grey lines, all other P25 samples of the same tissue class; purple line, position of c.1187G>A; dashed line, promoter/CpG island. **b**, Whole-gene *MUTYH* expression (transcripts per million, TPM) in every P25 bulk RNA-sequencing sample (*n* = 356), ordered by donor; open circles, non-carrier donors; filled symbols, carriers (squares, SMHT020; triangles, SMHT031; diamonds, SMHT043). **c**, TPM in all P25 samples (*n* = 356), carrier samples (*n* = 29: SMHT020, 4; SMHT031, 18; SMHT043, 7) and non-carrier samples (*n* = 327).

These observations extend the SBS18 phenotype from tumor to non-tumor tissue and from biallelic to monoallelic *MUTYH*. Recent tumor only studies have attributed SBS18 in carriers to haploinsufficiency^15^, showing cancers in which one allele is lost entirely. Our observation shows a single missense allele in normal tissue and interprets the SMHT043 phenotype as the consequence of a reduced-function (hypomorphic) allele.

The detection of a mutational phenotype consistent with oxidative DNA damage in non-neoplastic tissue supports models of pre-neoplastic activity previously inferred from tumors^15^. Detection in only one of three *MUTYH* heterozygotes carrying the same allele, however, suggests interindividual variability, as expected for an intermediate-activity allele whose effect may depend on additional modifiers^20^.

Pathogenic *MUTYH* alleles are carried by roughly 1 in 100 individuals, among the most common single-gene cancer-predispositions, yet carrier status is treated as a binary label and only a minority of carriers actually develop cancer^12^. A quantitative, tissue-level measure of accumulated oxidative mutational burden could separate molecularly active carriers such as SMHT043, who might warrant additional colorectal surveillance, from molecularly silent carriers such as SMHT020 and SMHT031. Because the signal is present in normal tissues, it offers a window in which the highest-risk carriers could be identified while intervention is still preventive.

More broadly, carrier status does not imply a neutral biological state. Heterozygous pathogenic variants in *ATM, BRCA2* and *PALB2* confer breast cancer susceptibility^21^, while *GBA1* heterozygotes have increased Parkinson disease risk^22^. Similarly, *SERPINA1* PiMZ heterozygosity increases susceptibility to chronic obstructive pulmonary disease, particularly in smokers^23^, and *CFTR* carriers have increased risks of several cystic fibrosis-related conditions^24^. These examples argue for systematic investigation of carrier phenotypes rather than the assumption that an unreported effect is absent. Multi-tissue resources such as SMaHT^16^ now make it possible to test directly whether heterozygous DNA-repair variants leave detectable mutational changes in normal (pre-neoplastic) tissues, as our *MUTYH* observation shows.

## Methods

### Cohort

The SMaHT P25 production cohort comprises the first 25 donors from the Somatic Mosaicism across Human Tissues Network^16^ (16 male, 9 female; mean age 68.8 years, median 66, range 42–89). A total of 381 tissue samples were analyzed (mean 15.2 per donor; range 11-19). Donor tissues were collected postmortem under the SMaHT Network consent^16^.

### Sequencing and variant calling

Tissues underwent Illumina short-read whole-genome sequencing (WGS) at the SMaHT Genome Characterization Centers (GCCs), and aligned CRAM files were obtained from the SMaHT data portal^16^. Somatic single-nucleotide variants were taken from the SMaHT Data Analysis Center (DAC) consensus tissue-level call set (SmahtSNV 2.0.0, filtered_calls_v2; GRCh38), one variant call format (VCF) file per tissue sample; only single-nucleotide variants with FILTER = PASS (EvidenceScore ≥ 1) were analyzed (536,748 sSNVs across the 381 samples; median 209 per sample). Germline variants were obtained from two sources: DAC consensus phased donor-level VCFs (release 2026.02.09), in which the individual genotype is derived from pooled read depth across all sequenced tissues of a donor; and independent calls made at the Baylor College of Medicine Human Genome Sequencing Center from the whole-blood WGS CRAM of each donor with Illumina DRAGEN v4.3.6. The *MUTYH* variant was present in both call sets in all three carriers.

### Germline DDR variant screen

Germline variants were screened against a curated set of 211 DNA damage repair genes assembled from the Wood laboratory human DNA-repair gene list^25^, the DDR compilation of He et al.^26^ and the DNA-damage-response gene set of Oak et al.^27^. Candidates were annotated against ClinVar by exact match on chromosome, position, reference and alternate allele (GRCh38) and stratified by pathogenicity evidence and expected mutational-signature family. The three pathogenic/likely pathogenic base-excision-repair candidates were the same allele, *MUTYH* c.1187G>A (p.Gly396Asp; rs36053993; ClinVar VCV000005294; chr1:45,331,556 C>T; NM_001128425 numbering; the same allele is p.Gly368Asp in NM_001048174 and p.Gly382Asp in the original report^5^). Carriers were heterozygous (0/1) by DRAGEN whole-blood genotype, and the variant was confirmed by inspection in the Integrative Genomics Viewer (IGV) and by Sanger sequencing (Fig. 1a,b).

### Mutational signature analysis

Per-sample SBS96 matrices were generated for all 381 tissue VCFs with SigProfilerMatrixGenerator^28^ (v1.3.6; GRCh38), and signatures were attributed against the COSMIC v3.4 SBS (GRCh38) reference with three methods. SigProfilerAssignment^18^ (v1.1.3) was run with cosmic_fit and default parameters against the full catalogue. MuSiCal^19^(v1.0.0) was run with the likelihood-bidirectional refit (threshold 0.001). SigFormer^29^ (v6 checkpoint SigFormer_v6_epoch300; code tag SigFormer_V6_no_OOC; simplex entmax15 output) was applied to the same matrices. For MuSiCal and SigFormer, each sample was fitted against a tissue-specific candidate subset of 14-17 COSMIC v3.4 signatures, formed as the union of a tissue-specific list of 11-12 signatures and a core set common to all tissues (SBS1, SBS3, SBS4, SBS5, SBS8, SBS13, SBS16, SBS17b, SBS18, SBS36, SBS40a, SBS40b, SBS89 and SBS92; SBS2 excluded from skin), as implemented in the SigFormer analysis pipeline (Extended Data Table 1); fitting against the full catalogue gave the same donor ranking (SMHT043 first with both methods). Per-donor SBS18 was summarized as the mean of per-sample SBS18 fractions across all tissues of the donor, and donors were ranked within each method; a tissue was considered SBS18-positive at ≥5% relative exposure. SBS36, which is in the candidate set of every tissue, was not assigned to any sample by SigProfilerAssignment. MuSiCal assigned 12-13 SBS36-attributed substitutions in total to each of SMHT020 and SMHT043 and none to SMHT031, fewer than to five non-carrier donors. SigFormer assigned a small SBS36 component to 76% of samples in the cohort (median of donor means 2.4%; range 0.9-5.9%); the donor mean was highest in SMHT043 (5.9%), with SMHT020 at 3.4% and SMHT031 at 1.8%.

### Statistics

Enrichment of SBS18 in SMHT043 was tested with a tissue-stratified donor-label permutation test (100,000 permutations; donor labels shuffled within each tissue stratum; per-sample SBS18 fractions; upper-tail P for the SMHT043 donor-level mean) and with a tissue-adjusted beta-binomial model of SBS18-attributed over total substitution counts per sample (SMHT043 indicator plus tissue type as fixed effects, with an overdispersion parameter; likelihood-ratio test against the tissue-only model; profile-likelihood 95% confidence interval). Both tests were run on the attributions of each method; the main text reports SigFormer, and SigProfilerAssignment and MuSiCal are shown in Extended Data Fig. 1. The enrichment was consistent when the logarithm of the per-sample sSNV count was added to the beta-binomial model as a covariate (odds ratios 2.16, 18.3 and 4.23 for SigFormer, SigProfilerAssignment and MuSiCal; likelihood-ratio *P* = 1.4 x 10^−6^, 7.5 x 10^−6^ and 4.5 x 10^−4^), and when the analysis was restricted to samples with ≥100 PASS sSNVs (358 samples, 16 from SMHT043; SigFormer permutation *P* = 1 x 10^−5^; odds ratio 1.87, 95% CI 1.40–2.42). *MUTYH* TPM in carrier versus non-carrier samples was compared with a two-sided Wilcoxon rank-sum test.

### Locus methylation

CpG methylation across *MUTYH* (chr1:45,329,242-45,340,925; GRCh38) was profiled from SMaHT long-read data. Per-read 5-methylcytosine (5mC) and 5-hydroxymethylcytosine (5hmC) calls from Oxford Nanopore (ONT), and combined 5mC+5hmC calls from ONT and PacBio, were summarized as the windowed percentage of methylated reads per sample. Each carrier sample was compared with all other P25 samples of the same tissue class (Fig. 2a); the digestive-tissue profiles are shown at larger scale, with the position of c.1187G>A and the promoter/CpG island marked, in Extended Data Fig. 2. Zoomed views of the promoter/CpG island (chr1:45,339,000-45,341,100) and a gene-body differentially methylated region (chr1:45,333,600-45,336,600) were also compared with samples of the same sub-tissue only, without median summarization. ONT calls were used as the primary measure.

### MUTYH expression

*MUTYH* expression was quantified from SMaHT bulk RNA-sequencing as whole-gene transcripts per million (TPM). Samples (*n* = 356; 29 from carriers) were analyzed pooled across tissues, with all P25 samples ordered by donor (Fig. 2b), and carrier samples were compared with non-carrier samples by two-sided Wilcoxon rank-sum test (Fig. 2c). Alignments at chr1:45,331,556 were inspected in IGV: the alternate allele was present in reads from every carrier tissue with RNA data and read coverage over exons downstream of the variant showed no reduction.

## Data availability

Whole-genome sequencing data, DAC consensus somatic (SmahtSNV 2.0.0) and germline variant calls, long-read methylation calls and bulk RNA-sequencing data for the SMaHT P25 cohort are available through the SMaHT Data Portal (https://data.smaht.org) under the Network’s controlled-access policy.

## Code availability

SigProfilerMatrixGenerator and SigProfilerAssignment are available at https://github.com/AlexandrovLab, MuSiCal at https://github.com/parklab/MuSiCal and SigFormer at https://github.com/zonglab/SigFormer.

## Acknowledgements

R.A.G. is partially supported by the NIH Somatic Mosaicism across Human Tissue (SMaHT) grant 1UM1DA058229.

## Author contributions

C.M.G. and R.A.G. conceived the study. C.M.G. performed the germline DDR screen, variant validation and mutational-signature analyses and wrote the manuscript. Y.Z. and C.Z. developed SigFormer and performed signature attribution and statistical analyses. Y.F. and F.J.S. performed long-read methylation analysis. L.F.P. performed RNA-sequencing analysis. E.V., S.B., Z.K. and J.F. performed sequencing, data QC, variant calling and data processing. M.-C.G. and H.D. supervised sample processing and sequencing. The SMaHT Consortium generated the P25 data.

R.A.G. supervised the study. All authors reviewed and approved the manuscript.

## Extended Data

**Extended Data Figure 1.**
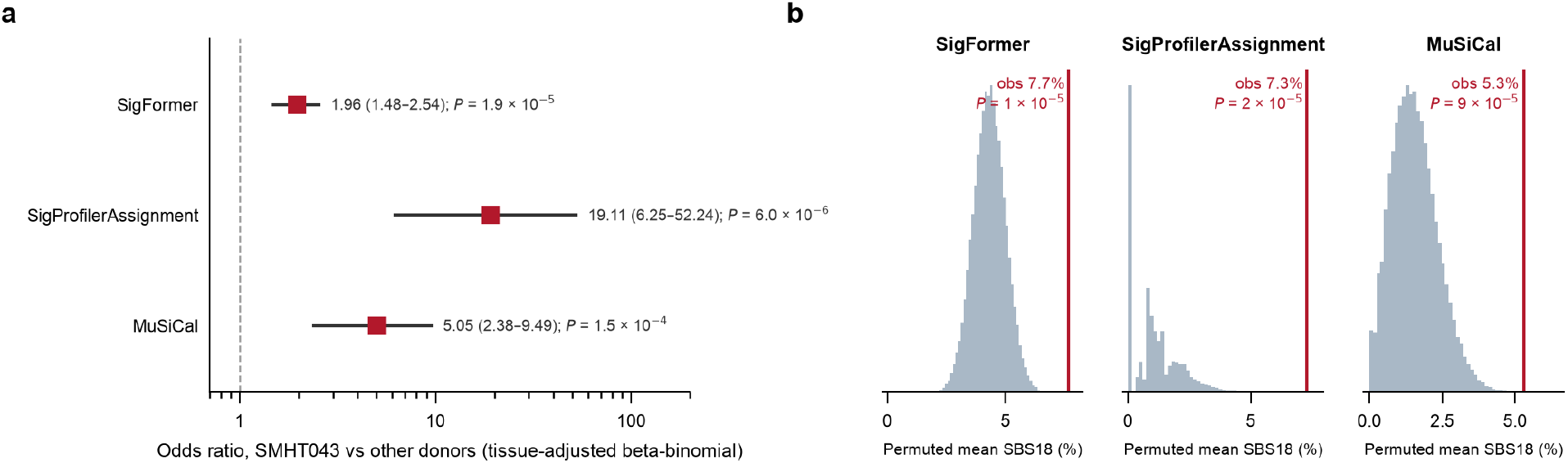
SBS18 enrichment in SMHT043 with each of the three attribution methods (SmahtSNV 2.0.0 PASS calls). **a**, SMHT043 odds ratio (square) with 95% profile-likelihood confidence interval from the tissue-adjusted beta-binomial model of SBS18-attributed over total substitutions, and the likelihood-ratio *P* value, for SigFormer (1.96, 1.48-2.54), SigProfilerAssignment (19.1, 6.2-52.2) and MuSiCal (5.05, 2.38–9.49); dashed line, odds ratio 1. **b**, Null distributions of the SMHT043 donor-level mean SBS18 fraction from 100,000 tissue-stratified donor-label permutations of each method’s attributions; red line, observed value; upper-tail *P* = 1 x 10^−5^ (SigFormer), 2 x 10^−5^ (SigProfilerAssignment) and 9 x 10^−5^ (MuSiCal).

**Extended Data Figure 2.**
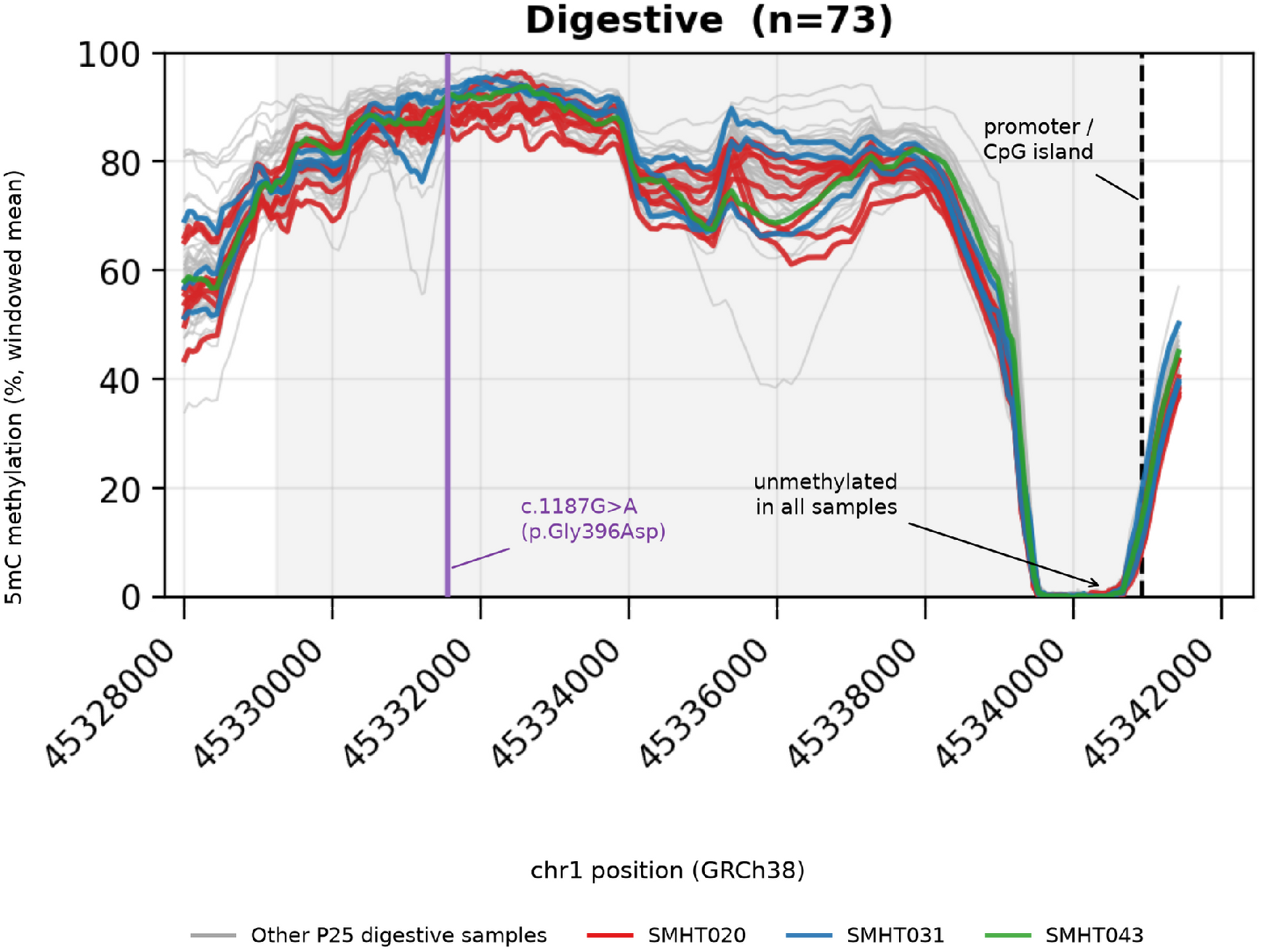
Long-read CpG methylation across *MUTYH* in digestive tissues, with the position of the carrier variant. Oxford Nanopore 5mC across *MUTYH* (chr1:45,328,000-45,342,000; GRCh38; gene body shaded) in all P25 digestive-tissue samples (*n* = 73; grey) and in the three carriers (SMHT020, red; SMHT031, blue; SMHT043, green), shown as the windowed mean percentage of methylated reads per sample. Purple line, position of c.1187G>A (p.Gly396Asp); dashed line, promoter/CpG island, which is unmethylated in every sample. The carriers track the cohort along the gene body, and the variant lies within a uniformly methylated gene-body region.

**Extended Data Table 1.** Tissue-specific candidate-signature subsets used for MuSiCal and SigFormer. For each of the 19 tissue types in the P25 cohort, the COSMIC v3.4 SBS signatures against which samples of that tissue were fitted by MuSiCal and SigFormer (14-17 per tissue: a tissue-specific list plus a core set common to all tissues, with SBS2 excluded from skin), and the number of samples per tissue. SigProfilerAssignment was fitted against the full catalogue.

| Tissue | n samples | Fitted candidate set (COSMIC v3.4) | n signatures fitted | Note |
|---|---|---|---|---|
| Abd-Aorta | 23 | SBS1, SBS2, SBS3, SBS4, SBS5, SBS8, SBS13, SBS16, SBS17b, SBS18, SBS36, SBS40a, SBS40b, SBS89, SBS92 | 15 |  |
| Adrenal | 19 | SBS1, SBS2, SBS3, SBS4, SBS5, SBS8, SBS13, SBS16, SBS17b, SBS18, SBS36, SBS40a, SBS40b, SBS89, SBS92 | 15 |  |
| Blood | 25 | SBS1, SBS2, SBS3, SBS4, SBS5, SBS8, SBS9, SBS13, SBS16, SBS17b, SBS18, SBS36, SBS40a, SBS40b, SBS89, SBS92 | 16 |  |
| Brain-Cereb | 15 | SBS1, SBS2, SBS3, SBS4, SBS5, SBS8, SBS13, SBS16, SBS17b, SBS18, SBS36, SBS40a, SBS40b, SBS89, SBS92 | 15 |  |
| Brain-Front | 15 | SBS1, SBS2, SBS3, SBS4, SBS5, SBS8, SBS13, SBS16, SBS17b, SBS18, SBS36, SBS40a, SBS40b, SBS89, SBS92 | 15 |  |
| Brain-Hippo | 24 | SBS1, SBS2, SBS3, SBS4, SBS5, SBS8, SBS13, SBS16, SBS17b, SBS18, SBS36, SBS40a, SBS40b, SBS89, SBS92 | 15 |  |
| Brain-Tempo | 15 | SBS1, SBS2, SBS3, SBS4, SBS5, SBS8, SBS13, SBS16, SBS17b, SBS18, SBS36, SBS40a, SBS40b, SBS89, SBS92 | 15 |  |
| Buccal_Swab | 9 | SBS1, SBS2, SBS3, SBS4, SBS5, SBS8, SBS13, SBS16, SBS17b, SBS18, SBS36, SBS40a, SBS40b, SBS89, SBS92 | 15 |  |
| Colon-Asc | 25 | SBS1, SBS2, SBS3, SBS4, SBS5, SBS8, SBS13, SBS16, SBS17b, SBS18, SBS36, SBS40a, SBS40b, SBS89, SBS92 | 15 |  |
| Colon-Desc | 24 | SBS1, SBS2, SBS3, SBS4, SBS5, SBS8, SBS13, SBS16, SBS17b, SBS18, SBS36, SBS40a, SBS40b, SBS89, SBS92 | 15 |  |
| Esophagus | 24 | SBS1, SBS2, SBS3, SBS4, SBS5, SBS8, SBS13, SBS16, SBS17b, SBS18, SBS36, SBS40a, SBS40b, SBS89, SBS92 | 15 |  |
| Heart | 22 | SBS1, SBS2, SBS3, SBS4, SBS5, SBS8, SBS13, SBS16, SBS17b, SBS18, SBS36, SBS40a, SBS40b, SBS89, SBS92 | 15 |  |
| Liver | 25 | SBS1, SBS3, SBS4, SBS5, SBS8, SBS13, SBS16, SBS17b, SBS18, SBS24, SBS36, SBS40a, SBS40b, SBS89, SBS92 | 15 | SBS22 not in COSMIC v3.4 (dropped) |
| Lung | 24 | SBS1, SBS2, SBS3, SBS4, SBS5, SBS8, SBS13, SBS16, SBS17b, SBS18, SBS29, SBS36, SBS40a, SBS40b, SBS89, SBS92 | 16 |  |
| Muscle | 23 | SBS1, SBS2, SBS3, SBS4, SBS5, SBS8, SBS13, SBS16, SBS17b, SBS18, SBS36, SBS40a, SBS40b, SBS89, SBS92 | 15 |  |
| Ovary | 4 | SBS1, SBS3, SBS4, SBS5, SBS8, SBS13, SBS16, SBS17b, SBS18, SBS36, SBS40a, SBS40b, SBS89, SBS92 | 14 |  |
| Skin-Abd | 25 | SBS1, SBS3, SBS4, SBS5, SBS7a, SBS7b, SBS7d, SBS8, SBS13, SBS16, SBS17b, SBS18, SBS36, SBS40a, SBS40b, SBS89, SBS92 | 17 | SBS2 excluded (skin) |
| Skin-Calf | 24 | SBS1, SBS3, SBS4, SBS5, SBS7a, SBS7b, SBS7d, SBS8, SBS13, SBS16, SBS17b, SBS18, SBS36, SBS40a, SBS40b, SBS89, SBS92 | 17 | SBS2 excluded (skin) |
| Testis | 16 | SBS1, SBS2, SBS3, SBS4, SBS5, SBS8, SBS13, SBS16, SBS17b, SBS18, SBS36, SBS40a, SBS40b, SBS89, SBS92 | 15 |  |

